# A Genetic Tool for Specific Tracking of Mature Neutrophils

**DOI:** 10.64898/2026.03.10.710957

**Authors:** Jiayu Cao, Hui Ping Yaw, Shilin Yi, Yanxi Zhou, Shifan Qin, Yuanyuan Wang, Raul da Costa, Lei Zhang, Dandan Wu, Changbin Chen, Melissa Ng, Immanuel Kwok, Oliver Soehnlein, Xiaoxiang Chen, Jieqing Wan, Lai Guan Ng

## Abstract

Tracking mature neutrophils remains challenging due to the lack of reliable cell surface markers. Although CD101 is a promising candidate for mature neutrophils, its stability under pathological conditions is unclear. Using a CD101-tdTomato reporter mouse model, we confirmed that the reporting system does not alter CD101 expression, and tdTomato fluorescence is predominantly expressed in mature neutrophils across peripheral tissues. Further analysis revealed that CD101^+^ and tdTomato^+^ neutrophils display identical characteristics of mature neutrophil, including poly-segmented nuclei, cell size, and key functions under homeostasis. By comparing tdTomato fluorescence with CD101 protein levels, we demonstrate that reduced CD101 expression under pathological states was not attributed to shedding or degradation. Our finding enhances CD101 as a robust and reliable marker of neutrophil maturity, providing a foundation for future applications in spatial transcriptomics and lineage tracing studies to dissect neutrophil heterogeneity and function.

**Highlights of the study:**

- In CD101-tdTomato homozygous mice, tdTomato is predominantly expressed in neutrophils and labels nearly 100% of mature neutrophils, aligning with the phenotype of CD101^+^ mature neutrophils;
- The CD101-tdTomato reporting system does not interrupt CD101 expression or neutrophil functions;
- CD101 remains a stable and reliable cell surface marker for labeling mature neutrophils, even under pathological conditions.

## Introduction

Neutrophils serve as the frontline defenders of the innate immune system. These cells rapidly migrate to sites of infection or injury, where they engulf pathogens, release antimicrobial molecules, and produce reactive oxygen species (ROS) to neutralize threats^1^. Their quick response is vital for host defense, playing a key role in combating bacterial infections, resolving tissue damage, and modulating immune responses. Despite their critical role in immunity, the heterogeneity of neutrophil populations—particularly the distinction between mature and immature subpopulations—has long posed a challenge to researchers. Neutrophil maturation is a complex process marked by intricate changes in morphology, function, and surface marker expression. These dynamic changes are often heterogeneous, making it difficult to clearly define distinct developmental stages and to track neutrophil behavior under both normal and diseased conditions^2^. The lack of specific, reliable markers complicates efforts to study neutrophil maturation and function, particularly in complex tissue environments^3^. Traditionally, expression levels of markers such as Ly6G and CD11b have been used to assess neutrophil maturation, however, this method remains inherently subjective^4, 5^. Furthermore, CXCR2 has been used to distinguish between mature and immature subpopulations, but its expression is highly context-dependent and influenced by activation states, making it an imprecise marker for defining neutrophil maturity^6^.

CD101, a cell surface glycoprotein, has emerged as a reliable marker of neutrophil maturation, characterized by high expression on mature neutrophils and minimal or undetectable expression on immature and early precursors^7^. This marker thus provides a valuable tool to refine our understanding of neutrophil development and function. However, tracking CD101^+^ neutrophils in tissues using traditional staining methods is difficult due to technical limitations, such as antibody penetration or tissue fixation issues^8^. The complexity of tissue microenvironments and the limitations of immunohistochemistry often hinder precise identification of mature neutrophils^9^.

Another critical question is whether CD101 remains a stable marker under challenged conditions. Neutrophils undergo rapid phenotypic and functional changes during stress, including upregulation of activation markers, release of proteases, and alterations in surface protein expression^10^. Like many surface proteins, CD101 may be subjected to regulation through mechanisms such as proteolytic shedding, internalization, or degradation, potentially compromising its reliability as a marker of neutrophil maturation under inflammatory or pathological conditions. A well-known example is the shedding of surface markers such as CD62L during neutrophil activation, highlighting the need for caution when using surface markers to define cell maturation or activation stages^12^. Similarly, a decrease in CD101^+^ cells may indicate changes in neutrophil populations, such as their mobilization from bone marrow to peripheral tissues^13^. Therefore, it’s important to validate the reliability of CD101 as a surface marker to distinguish between a true reduction in cell numbers and a stress-induced downregulation of the marker.

To evaluate this further, we developed a CD101-tdTomato reporter mouse model. In this mouse model, the CD101-tdTomato reporting system shows synchronized expression of CD101 and tdTomato, with tdTomato labelling nearly all of the CD101^+^ mature neutrophils across tissues. We confirmed that CD101^+^ and tdTomato^+^ mature neutrophils share identical nucleus morphology and function traits. Moreover, tdTomato labeling efficiency remains unchanged under stress conditions. Our results validate CD101 as a reliable cell surface marker and provide a powerful tool for investigating neutrophil heterogeneity, thereby enabling more precise studies of neutrophil function in immunity and disease.

## Result

### Consistent Expression of CD101 and tdTomato Across Tissues in Homeostasis

CD101 is highly expressed by mature neutrophil^7^. In order to investigate CD101 as a stable cell surface marker, we generated a mouse line in which the exon of CD101 is knocked in an allele encoding Cre recombinase and the fluorescent protein tdTomato separated by a self-splicing T2A peptide (**Fig. 1A**). Gene-targeted animals were verified by PCR (**Supplementary Fig.1A**). CD101 expression levels in bone marrow (BM) neutrophils were similar across wild-type, heterozygous, and homozygous CD101-tdTomato mice, indicating that the genetic modification did not alter CD101 gene regulation or protein expression (**Fig. 1B, C and Supplementary Fig. 1B**). Moreover, the homozygous mice show high percentage of tdTomato^+^ cells compared to heterozygous mice, aligning closely with CD101^+^ population. This observation extend beyond BM, with similar patterns of CD101 consistency and enriched tdTomato expression observed in the spleen and blood of homozygous mice (**Fig. 1B, C**). Correlation analysis revealed a strong positive correlation between CD101 and tdTomato expression in BM, blood, and spleen of homozygous mice (**Fig. 1D and Supplementary Fig. 1C**). In CD101-tdTomato homozygous mice, tdTomato expression was predominantly restricted to Ly6G^+^ neutrophils, demonstrating the specificity of the CD101-tdTomato system for labelling neutrophils compared to heterozygous mice (**Fig. 1E and Supplementary Fig. 1D**). Furthermore, tdTomato labeled nearly 100% of CD101^+^ cells in BM, blood, and spleen, indicating high labeling efficiency (**Fig. 1F**). Taken together, these findings collectively demonstrate that the CD101-tdTomato homozygous mouse model is a highly specific and reliable tool for tracking mature neutrophils across various tissues under physiological conditions.

**Figure 1.**
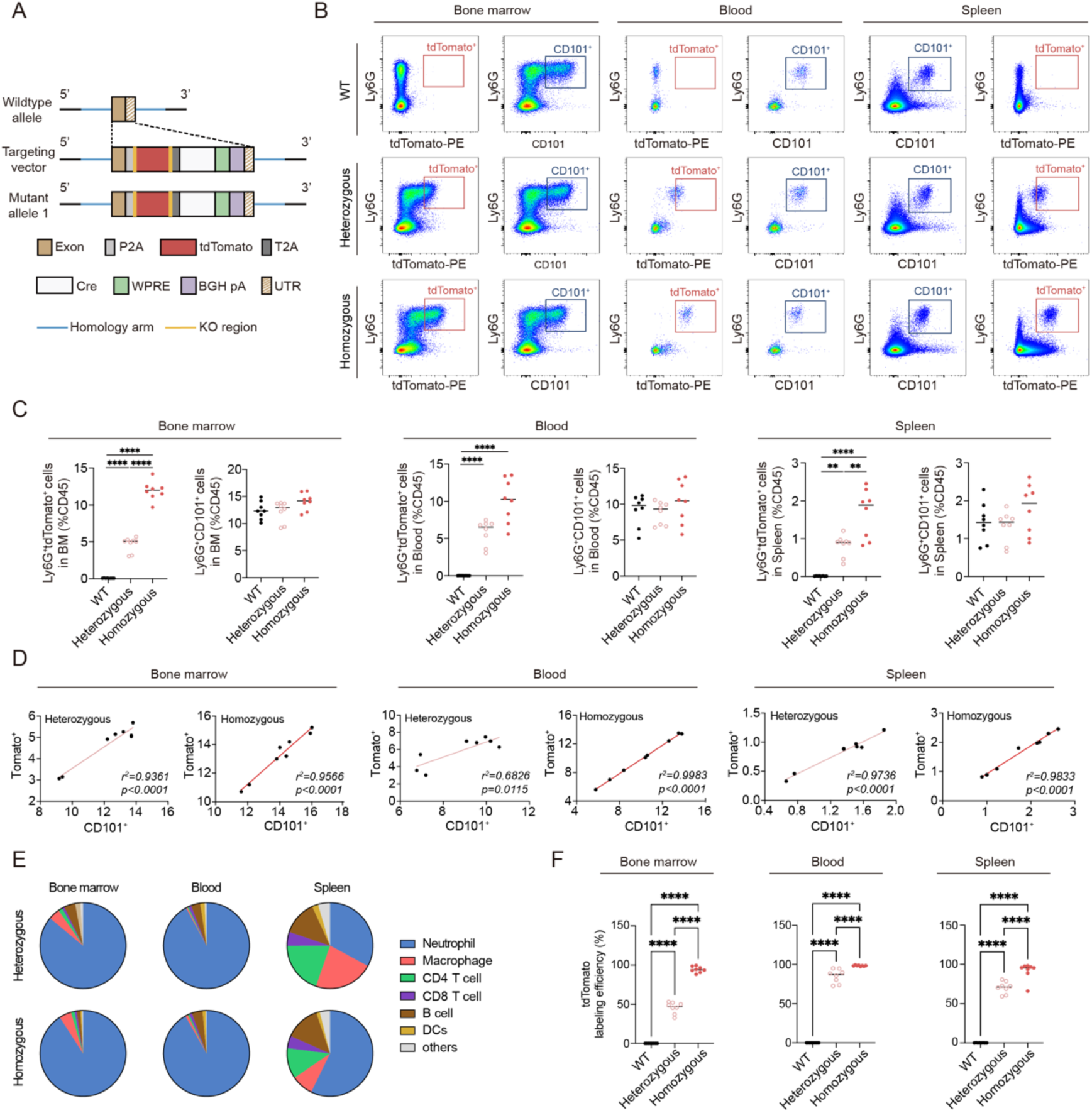
CD101 and tdTomato expression in neutrophil in peripheral tissues at the steady states. **A.** Targeting strategy for the generation of CD101-tdTomato mice; **B.** Representative flow cytometry plots showing tdTomato^+^ and CD101^+^neutrophil population in bone marrow (BM), peripheral blood, and spleen from wild type (WT), heterozygous, and homozygous CD101-tdTomato mice; **C.** Quantification of the proportion of tdTomato^+^ and CD101^+^ neutrophils in peripheral organs (ANOVA, *n* = 8 per group); **D.** Correlation analysis of tdTomato^+^ and CD101^+^ neutrophil populations in bone marrow, peripheral blood, and spleen of heterozygous and homozygous mice, with Pearson correlation coefficients and *p*-values indicated (*n* = 8 per group); **E.** tdTomato expression profiles in immune cell subsets from the bone marrow, peripheral blood, and spleen of heterozygous and homozygous mice (*n* = 8 per group); **F.** Statistical analysis of tdTomato labeling efficiency in bone marrow, peripheral blood, and spleen of heterozygous and homozygous mice (ANOVA, *n* = 8 per group).

### CD101^+^ and tdTomato^+^ Neutrophils Exhibit Mature Nuclear Morphology Across Tissues

To assess the reliability of tdTomato and CD101 as markers for distinguishing mature from immature neutrophil, we checked the nucleus morphology by Giemsa staining^1^ (**Supplementary Fig. 2**). In the BM of both homozygous CD101-tdTomato and WT mice, tdTomato^+^ and CD101^+^ neutrophils showed the characteristic poly-segmented nuclei of mature neutrophils, while tdTomato^-^ and CD101^-^ cells exhibited a typical donut-shaped nucleus of immature neutrophils. This distinction was consistently observed in blood and spleen, where tdTomato^+^ and CD101^+^ cells consistently displayed poly-segmented nuclei (**Fig. 2A**), indicating tdTomato and CD101 reliably identify mature neutrophils across different tissues. Statistical analysis of tdTomato^+^ cells revealed that nearly all were mature neutrophils, supporting tdTomato as a specific marker for neutrophil maturity across tissues (**Fig. 2B**). Measurements of tdTomato^+^ and CD101^+^ cells indicated a consistent diameter of approximately 15 μm for mature neutrophils in multiple peripheral tissues, compared to about 10 μm for immature (tdTomato^-^ / CD101^-^) cells (**Fig. 2C**).

**Figure 2.**
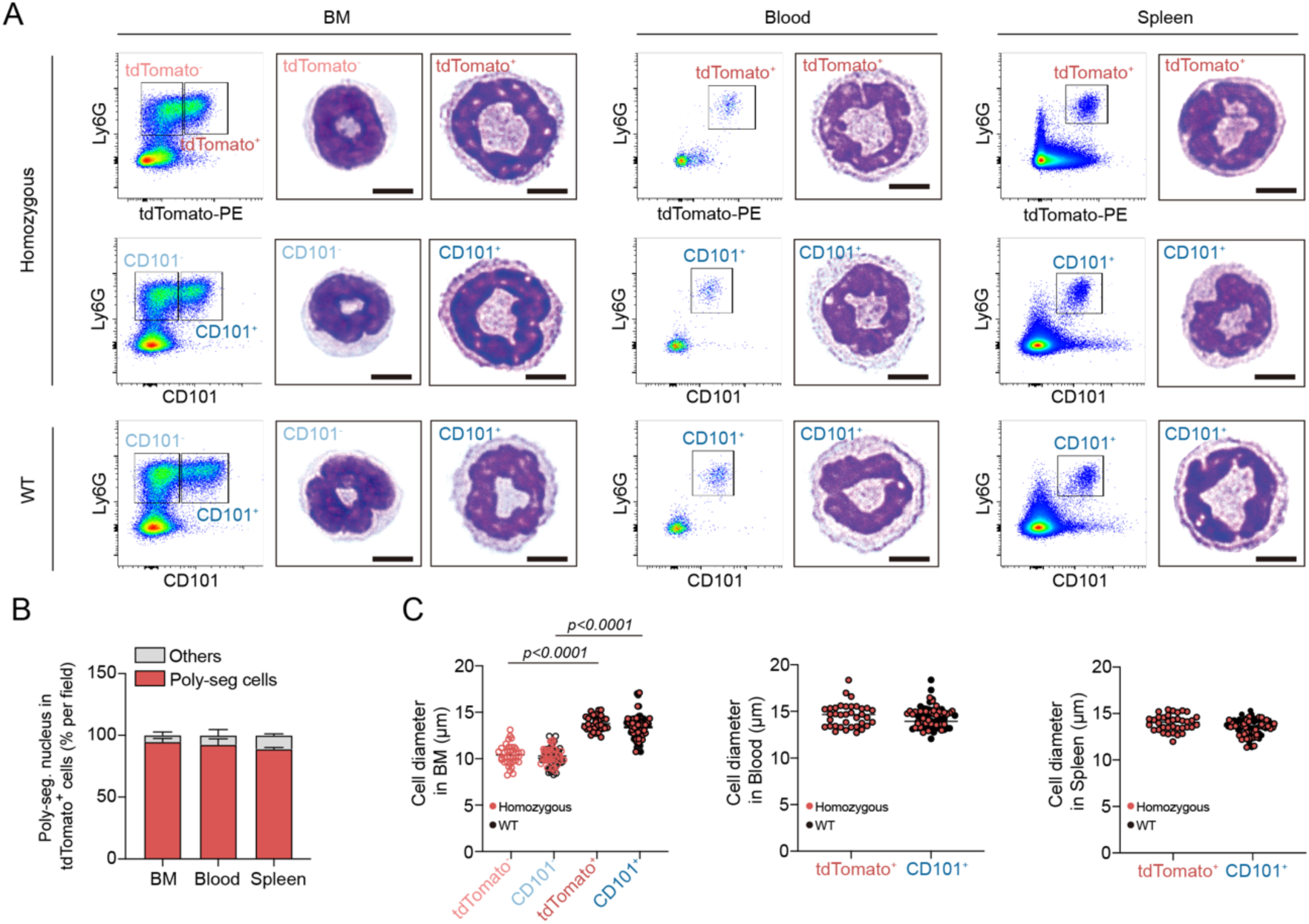
Mature morphological features of CD101^+^ and tdTomato^+^ neutrophils across tissues. **A.** Representative sorting plots and corresponding Giemsa staining images of sorted neutrophils from BM, blood, and spleen in homozygous CD101-tdTomato and WT mice (Scale bar = 5μm); **B**. Quantification of the proportion of poly-segmented nuclei in tdTomato^+^ sorted neutrophils from BM, peripheral blood, and spleen of homozygous and WT mice (ANOVA, *n* = 5 per group); **C**. Statistical analysis of cell diameter in sorted neutrophils from BM (left), peripheral blood (middle), and spleen (right) of homozygous and WT mice (Two-way ANOVA in BM, U-test in blood and spleen, *n* = 36 per group).

### Functional Characterization of CD101^+^ and tdTomato^+^ Mature Neutrophils

To determine whether the genetic modification in the CD101-tdTomato mouse model affects neutrophil physiology, we first compared the viability of neutrophils from CD101-tdTomato mice with that of WT controls. We observed a gradual increase in cell death and apoptosis over time, with identical trends for CD101^-^ and tdTomato^-^ cells, indicating consistent behavior in the immature population (**Fig. 3A and Supplementary Fig. 3A, B**). In contrast, mature neutrophils (CD101^+^ / tdTomato^+^) displayed earlier apoptosis and cell death compared to immature neutrophils, with synchronized changes in CD101^+^ and tdTomato^+^ cells (**Fig. 3B and Supplementary Fig. 3C**). This highlights distinct survival kinetics between mature and immature neutrophils, with CD101 and tdTomato reliably distinguishing these populations. Furthermore, we examined reactive oxygen species (ROS) production, a key effector mechanism critical for neutrophil-mediated hose defense^1^. To determine if neutrophils from WT and homozygous CD101-tdTomato mice produce similar ROS levels, we stimulated neutrophils with phorbol myristate acetate (PMA). Neutrophils from bone marrow and spleen between strains showed comparable ROS production, suggesting that the CD101-tdTomato reporter system did not affect this critical function (**Fig. 3C, D and Supplementary Fig. 3D**). Another key biological feature of neutrophils is their migratory capacity. To assess chemotaxis, a vital neutrophil function for their rapid migration to infection sites^2^, we FACS-sorted CD101^+^ / tdTomato^+^ neutrophils from bone marrow and performed a transwell migration assay. CD101^+^ and tdTomato^+^ mature neutrophils exhibited nearly identical chemotaxis capacity, while CD101^-^ / tdTomato^-^ immature neutrophils showed similar but distinct performance, indicating a functional difference linked to maturity (**Fig. 3E**).

**Figure 3.**
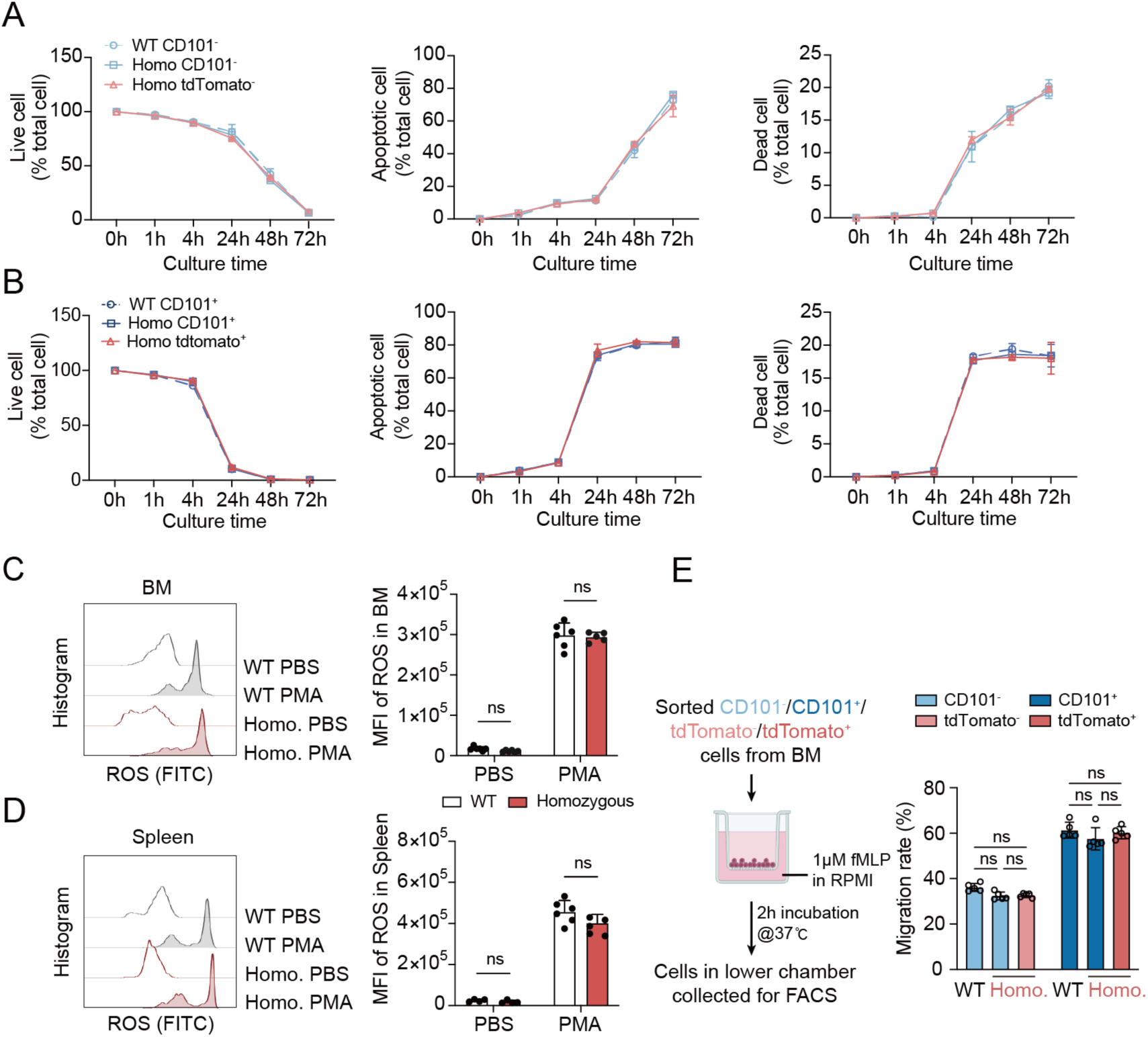
Functional assessment of CD101^+^ and tdTomato^+^ mature neutrophils. **A.** Quantification of live cell (left), apoptotic cell (middle), and dead cell (right) at various time points in culture for sorted immature CD101^-^ and tdTomato^-^ neutrophils from BM of WT and homozygous mice (*n* = 6 per group, 3 independent experiments); **B.** Quantification of cell viability (left), apoptosis (middle), and cell death (right) at various time points in culture for sorted mature CD101^+^ and tdTomato^+^ neutrophils from BM of WT and homozygous mice (*n* = 6 per group, 3 independent experiments); **C**, **D.** Representative flow cytometry histograms (left) and corresponding quantification (right) of ROS production in Ly6G^+^ neutrophils sorted from BM and spleen of WT and homozygous CD101-tdTomato mice upon phorbol PMA stimulation (U-test, *n* = 6 per group, 3 independent experiments); **E.** Schematic diagram of the neutrophil transwell migration assay (left) and quantification of migration rates for sorted WT and homozygous neutrophil populations from BM (Two-way ANOVA, *n* = 6 per group, 3 independent experiments).

### CD101 and tdTomato as Stable Markers in Neutrophil Stress Responses

To further determine whether CD101 is a stable cell surface marker under stress conditions, we evaluated CD101 and tdTomato expression in CD101-tdTomato mice following lipopolysaccharide (LPS) infection (**Supplementary Fig. 4A**). After LPS administration, the proportion of CD101^+^ and tdTomato^+^ mature neutrophils decreased in the BM of WT and homozygous CD101-tdTomato mice. Both CD101 and tdTomato signals were diminished to a comparable extent, with a high correlation between their expression levels in BM; notably, the efficiency of CD101 antibody labeling was preserved (**Fig. 4A-C and Supplementary Fig. 4B, C**). This observation indicates that CD101 is neither shed nor degraded, but rather that the actual frequency of CD101^+^ (tdTomato^+^) neutrophils is reduced. This pattern of decreased CD101^+^ and tdTomato^+^ cell proportions with stable labeling was also observed in spleen and blood, suggesting that LPS challenge reduces the mature neutrophil population without compromising CD101 integrity (**Fig. 4A-C and Supplementary Fig. 4B, C**).

**Figure 4.**
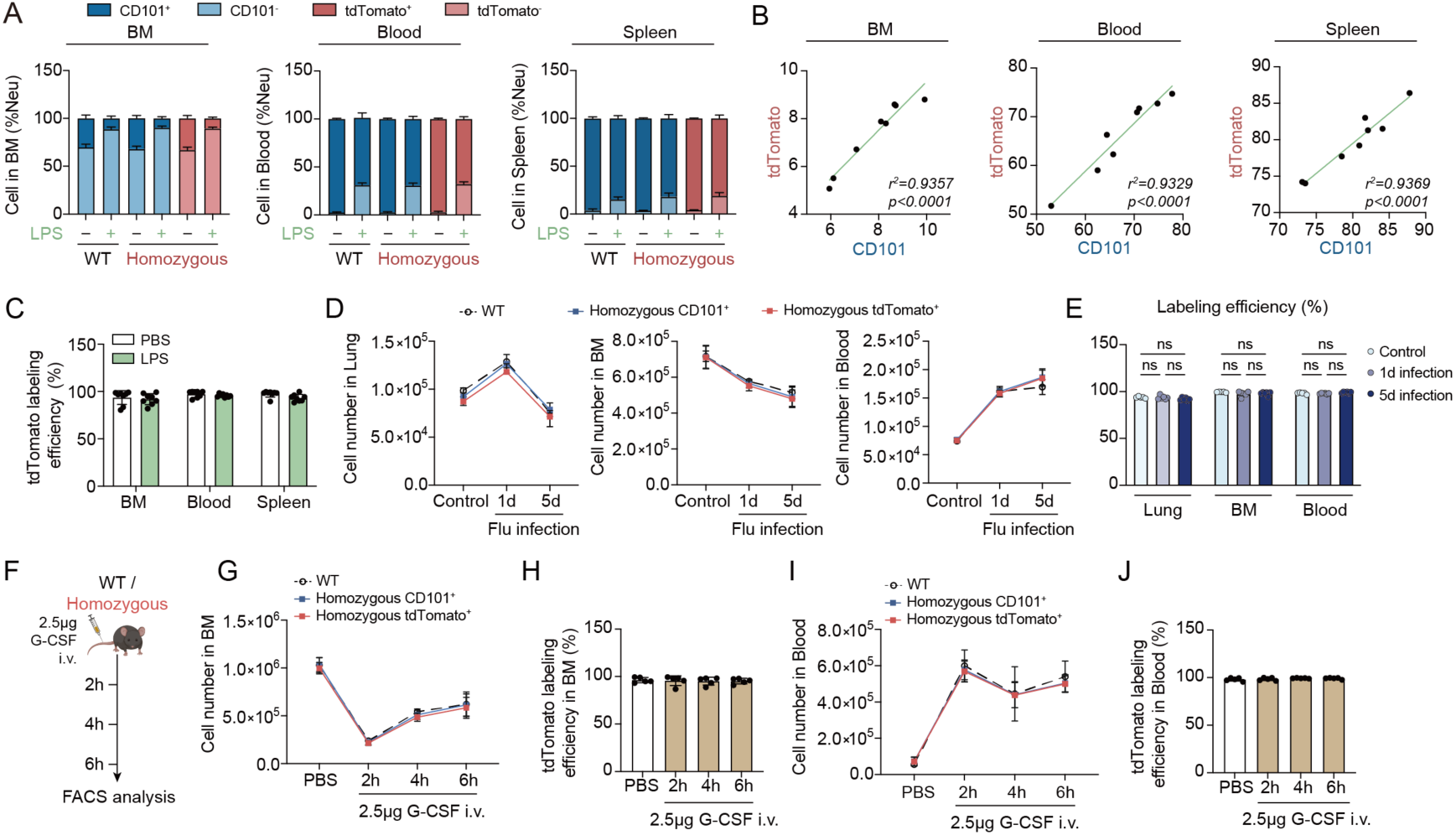
Stability of CD101 and tdTomato expression in neutrophils under stress conditions. **A.** Flow cytometry analysis showing changes in CD101^+^ and tdTomato^+^ mature neutrophil populations in BM (left), peripheral blood (middle), and spleen (right) of WT and homozygous CD101-tdTomato mice under control conditions versus LPS challenge ( *n* = 8 per group); **B.** Correlation analysis of tdTomato^+^ and CD101^+^ neutrophil populations in BM (left), peripheral blood (middle), and spleen (right) of CD101-tdTomato homozygous mice treated with LPS, with Pearson correlation coefficients and *p*-values indicated (*n* = 8 per group); **C.** Statistical analysis of tdTomato labeling efficiency in BM (left), peripheral blood (middle), and spleen (right) of homozygous mice under control and LPS-treated conditions (U-test, *n* = 8 per group); **D.** Quantification of CD101^+^ and tdTomato^+^ mature neutrophils in lung (left), BM (middle), and blood (right) of WT and homozygous mice at 1 day and 5 days after intratracheal injection of H1N1 virus. (*n* = 5-6 per group); **E.** Calculation of tdTomato labeling efficiency in lung, BM and blood under control conditions and at 1 day and 5 days after intratracheal injection (ANOVA, *n* = 5-6 per group); **F.** Schematic diagram illustrating the experimental design for G-CSF injection to simulate emergency granulopoiesis; **G.** Quantification of CD101^+^ and tdTomato^+^ mature neutrophil populations in BM of WT and homozygous mice at 2, 4, and 6 hours post-G-CSF injection (*n* = 5 per group); **H.** Statistical analysis of tdTomato labeling efficiency in BM of homozygous mice under control conditions and at different time points post-G-CSF injection (ANOVA, *n* = 5 per group); **I.** Quantification of CD101^+^ and tdTomato^+^ mature neutrophil populations in peripheral blood of WT and homozygous mice at 2, 4, and 6 hours post-G-CSF injection (*n* = 5 per group); **J.** Statistical analysis of tdTomato labeling efficiency in peripheral blood of homozygous mice under control conditions and at different time points post-G-CSF injection (ANOVA, *n* = 5 per group);

In order to further evaluate the stability of CD101 in a viral infection-induced stress environment, we performed intratracheal injections of Influenza A virus (A/Puerto Rico/8/1934, H1N1) into the WT and CD101-tdTomato reporter mice (Fig. 4D). Following infection, we observed a rapid increase in the infiltration of CD101^+^ and tdTomato^+^ mature neutrophils in the lung at 1day post-infection (dpi), which then gradually declined by 5 dpi (**Fig**. **4D** **left**). Concurrently, the number of these mature neutrophils in the BM showed a progressive decrease, while their frequency in the peripheral blood steadily rose, reflecting a systematic mobilization from the marrow to the site of infection (**Fig**. **4D** **middle and right**). Notably, the dynamic changes in CD101^+^ cell counts were highly consistent between WT and homozygous CD101-tdTomato reporter mice, suggesting that the genetic insertion does not interfere with the physiological recruitment or maturation of neutrophils. Furthermore, by calculating the labeling efficiency at both 1 and 5 dpi, we found that tdTomato successfully marked CD101^+^ cells throughout the course of infection in lung, BM and blood (**Fig**. **4E**). These findings reinforce the conclusion that CD101 is a robust and stable marker for mature neutrophils in peripheral organs, even under the intense inflammatory conditions of viral pneumonia.

To investigate the response of CD101-tdTomato reporter mice to emergency granulopoiesis, we intravenously injected 2.5 µg of G-CSF, a potent inducer of granulopoiesis, and analyzed neutrophil populations in blood and bone marrow dynamically (**Fig. 4F and Supplementary Fig. 5A**). 2 hours after G-CSF injection, bone marrow neutrophils in both WT and homozygous mice sharply declined, followed by partial recovery, reflecting rapid mobilization to peripheral tissues (**Supplementary Fig. 5B**). Mature neutrophils (CD101^+^ / tdTomato^+^) showed a similar sharp decline and slight recovery, with both populations displaying identical trends (**Fig. 4G**). The labeling efficiency remained above 90% throughout, confirming synchronized changes in CD101 and tdTomato expression (**Fig. 4H**). In contrast, blood neutrophils in WT and homozygous mice rapidly increased after G-CSF injection, remaining elevated above controls at 4 and 6 hours, with CD101^+^ and tdTomato^+^ mature neutrophils showing synchronized increases (**Fig. 4I, J and Supplementary Fig. 5B, C**). These findings demonstrate that the CD101-tdTomato system did not alter emergency granulopoiesis and reliably marks mature neutrophils with stable labeling under G-CSF-induced stress.

### Distinguishing Mature Neutrophils from Myeloid Cells

To enable identification of mature neutrophils from total myeloid cell population, we crossed homozygous CD101-tdTomatomice with Lysozyme-GFP mice, a well-established tool strain for labeling myeloid cells^19^. Flow cytometric analysis of these offspring revealed that tdTomato^+^ GFP^+^ population consistently overlapped with the CD101^+^ GFP^+^ phenotype, indicating that the tdTomato expression faithfully marks mature neutrophils. Importantly, the *CD101*-tdTomato x *LysM*-GFP mice exhibited clear separation of tdTomato^+^ mature neutrophils from the tdTomato^-^GFP^+^ population (comprising immature neutrophils, monocytes, and macrophages) across tissues/organs analyzed (**Fig. 5 and Supplementary Fig. 6**). These findings validate the compatibility of the CD101-tdTomato reporter with other fluorescent genetic tools and highlights its utility for high-resolution lineage analysis, providing a reliable method to visualize mature neutrophils distinct from other inflammatory myeloid cells.

**Figure 5.**
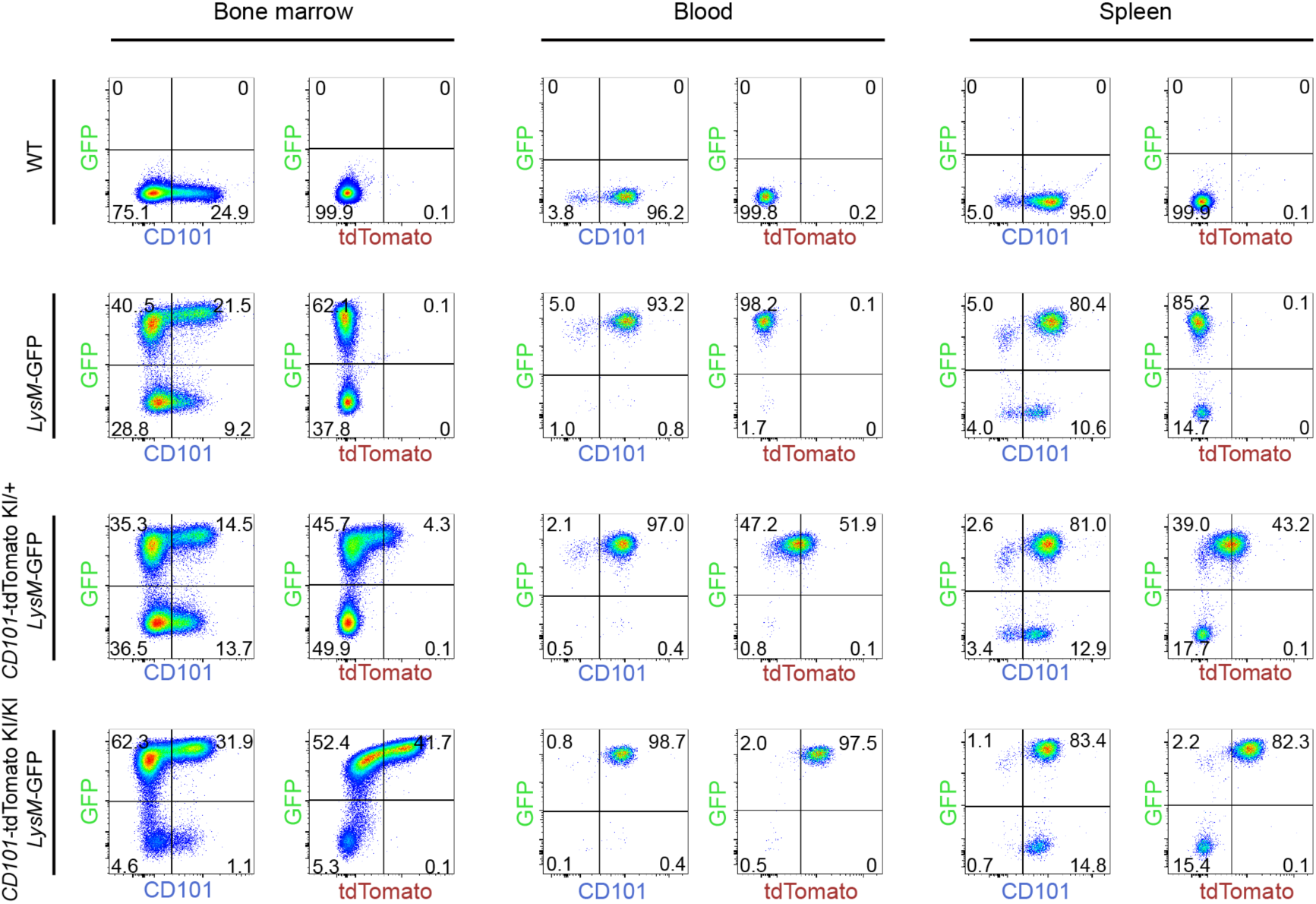
Tracking mature neutrophils in CD101-tdTomato Lysozyme GFP mice. Representative flow cytometry plots demonstrating the identification of GFP^+^ CD101^+^ and GFP^+^ tdTomato^+^ mature neutrophils within the total GFP^+^ myeloid cell population in bone marrow (left), peripheral blood (middle), and spleen (right) across four mouse strains: wild-type (WT; top row), *Lysozyme*-GFP (*LysM*-GFP, second row), heterozygous *CD101*-tdTomato × *LysM*-GFP (third row), and homozygous *CD101*-tdTomato × *LysM*-GFP (bottom row), with population frequencies indicated.

## Discussion

Neutrophils play a vital role in immunity, yet distinguishing mature neutrophils from their immature counterparts remains a challenge due to the absence of specific, reliable markers. This difficulty arises because neutrophil maturation involves complex changes in morphology, function, and surface marker expression, which are not always well-defined^3^. Reliable markers are essential for studying neutrophil biology, especially in understanding their roles in health and disease. CD101 has been reported as a promising candidate due to its predominant expression on mature neutrophils, offering greater precision than traditional markers, though its expression under stress remains unclear^7^. Our study introduces a novel CD101-tdTomato reporter mouse model to confirm that CD101^+^ and tdTomato^+^ neutrophils share hallmark traits of maturity, such as poly-segmented nuclei, larger cell size (∼15 μm), and enhanced functions like ROS production and chemotaxis migration. The model achieves nearly 100% labeling efficiency across bone marrow, blood, and spleen. Moreover, we showed that CD101 reduction under stress reflects population shifts rather than marker shedding or degradation. Furthermore, by crossing CD101-tdTomato mice with Lysozyme-GFP mice—a widely used strain for labeling general myeloid cells^19^—we successfully distinguished mature neutrophils (tdTomato^+^ GFP^+^) from general myeloid population (tdTomato^-^GFP^+^). Our core conclusion is that CD101 is a specific marker for mature neutrophils under homeostasis, with the CD101-tdTomato system providing a reliable tool for tracking these cells. The system preserves CD101 expression and neutrophil function, making it ideal for imaging and functional studies. Unlike antibody-based staining, which struggles with tissue penetration, our fluorescent reporter enables robust visualization of CD101^+^ neutrophils in complex tissues^8^. The high labeling efficiency of our reporter mouse model make it a significant improvement over other reporter systems, such as those for Ly6G, which are less specific to mature neutrophils^14^.

Compared to prior studies, our work provides a more comprehensive evaluation by testing CD101 under both steady-state and stress conditions, offering a clearer understanding of its behavior^6, 11^. Our findings have important implications for the use of CD101 as a marker of neutrophil maturity, particularly in disease settings where neutrophil phenotypes are altered. By establishing the conditions under which CD101 remains a faithful indicator of maturity, we lay the groundwork for its application in cutting-edge technologies like spatial transcriptomics and lineage tracing. Such tools could ultimately reveal the spatial organization and developmental origins of neutrophil subsets, deepening our understanding of their roles in immunity and disease. Moreover, our reporter mouse model offers a versatile platform for future studies of neutrophil biology, bridging the gap between traditional surface marker analysis and modern omics approaches. The CD101-tdTomato model also paves the avenue for high-resolution imaging of mature neutrophils in tissues, overcoming the limitations of immunohistochemistry, which often fails in dense microenvironments^9^. For example, intravital microscopy could leverage tdTomato fluorescence to track CD101^+^ neutrophils in real-time during infection or tissue injury, offering insights into their migration and effector functions^15^.

Despite its strengths, our study has limitations. We focused on stressed conditions (LPS, viral infection, and G-CSF), which may not fully represent CD101 behavior in chronic conditions like autoimmune diseases^16^. Additionally, our findings are based on murine systems, and the role of CD101 in human neutrophils requires validation, as species-specific differences in marker expression may exist^17^. Minor variations in tdTomato fluorescence intensity could also affect sensitivity in low-expression settings, potentially limiting detection in certain contexts. Future studies should test CD101 in chronic disease models, such as rheumatoid arthritis or tumor microenvironments, to assess its stability across diverse settings. Validating CD101 expression in human samples and explore its role in mature neutrophils across diverse human disease models to uncover its potential relevance for disease expression profiling and diagnostic applications. Combining the CD101-tdTomato model with advanced techniques like mass cytometry or spatial transcriptomics could further dissect neutrophil subsets, revealing their functional roles in complex diseases. Pairing CD101 with other markers, such as CXCR4, could enhance specificity for mature neutrophils in inflammatory contexts^18^. These steps would strengthen utility of CD101 and address its dynamic expression under stress.

In summary, our study shows the CD101-tdTomato model as a powerful tool for tracking mature neutrophils with high specificity. Its future applications in imaging and lineage tracing offer new avenues for studying neutrophil biology. While CD101 shows promise as a reliable marker, its dynamic expression during inflammation calls for further investigation. By providing a platform for precise, in vivo insights, this model sets the stage for deeper explorations of neutrophil roles in health and disease, potentially guiding the development of targeted therapies.

## Material and Methods

### Mice

All animal experiments were performed in accordance with the guidelines for the use of experimental animals and were approved by the local authorities in Shanghai Jiao Tong University School of Medicine (JUMC2023-161-A). For the experiments, mice of both sexes were used. Mice used were between 7 and 10 weeks old.

The Cd101-tdTomato-iCre knock-in mouse model (C57BL/6JCya-Cd101em1 (tdTomato-iCre) /Cya) was generated by Cyagen Biosciences (Suzhou) Inc. To generate this mice, the TGA stop codon of Cd101 were replaced with a cassette containing the open reading frame of tdTomato and codon optimized iCre recombinase separated by a self-splicing P2A and T2A peptide ,thereby using the endogenous promoter from Cd101 for the P2A-tdTomato-T2A-iCre expression. A polyadenylation signal (WPRE-BGH pA)was inserted at the 3’end of the P2A-tdTomato-T2A-iCre sequence in order to terminate transcription. The “P2A-tdTomato-T2A-iCre-WPRE-BGH pA” targeting vector was generated using BAC clones from the C57BL/6J RPCI-23 BAC library . The fertilized eggs were obtained from C57BL/6JCya wild-type mice. The Cas9 protein, gRNA (gRNA:5’-GACACTACTGAGGAGCACCTGGG-3’), and the targeting vector were co-injected into fertilized eggs. The injected embryos were cultured in KSOM medium overnight and those developed to the two-cell stage were transferred into the oviduct of pseudopregnant ICR female mice. The F0 founder mice were identified by PCR anaysis were identified by PCR and Sanger sequencing, which were bred to wild type mice to test germ line transmission and F1 animal generation. The genotype F1 mice was also confirmed by PCR.

### Mouse genotyping

Tips of mouse tails were incubated at 100 °C for 10 minutes in 200 μL of 50 mM NaOH solution. After incubation, 50 μL of 1 M Tris-HCl solution was added and the DNA mixture was vortexed thoroughly. The typical PCR sample consisted of a 20-μL volume containing 1 μL of the primers (10 μM) for mutant allele (forward: 5’-CTAGTTGCCAGCCATCTGTTGTTT-3’; reverse: 5’-ACTATCTGCTTCTGAAGGATGCAA-3’) or for wildtype allele (forward: 5’-TCTTCTGTGTTTCTCTGGTTGCAG-3’; reverse: 5’-CCAAGGGAATAGATGAGTGGGAGA-3’). Each reaction also contained 4 μL of DNA mixture, 4 μL of ddH_2_O and 10 μL of 2 × Taq Master Mix (Dye Plus) (Vazyme, Cat# P212). The following PCR conditions were applied: 3 min, 94 °C for initial denaturation; 30 s, 94 °C cyclic denaturation; 35 s, 60 °C cyclic annealing; 35 s, 72 °C cyclic elongation for a total of 35 cycles, followed by a 5-min 72 °C elongation step. The PCR result for mutant allele was 431 bp and for wildtype allele was 580 bp.

### Drug administrations

**LPS challenge.** Both WT and tdTomato homozygous mice received once injection of LPS (Sigma-Aldrich) dissolved in 1x PBS at a dose of 200 ng in a final volume of 100 μl. The control group was administrated 1x PBS injections at the same volume, routine and timing as the experimental group. Mice were sacrificed and peripheral organs were collected for further experiments 2 hours after LPS or 1x PBS injection.

**Virus-induced pneumonia model.** For the influenza infection model, WT and homozygous CD101-tdTomato reporter mice were anesthetized by isoflurane and administered 40μl of Influenza A virus (A/Puerto Rico/8/1934, H1N1) at a dose of 10xLD_50_ via intratracheal injection. Control group received an equal volume of sterile PBS. Mice were euthanized at 1 and 5 days post infection, and lung, peripheral blood and BM were then harvested and processed for further analysis.

**Emergency granulopoiesis.** The G-CSF injected to induce emergency granulopoiesis as previously characterized^1^. The stock solution (in DMSO) were dissolved in PBS and injected intravenously at a dose of 2.5 μg in a final volume of 100 μl. The control group was administrated control injections (PBS) at the same volume, routine and timing as for the intervention group. Mice were sacrificed at 2 hours, 4 hours and 6 hours after injection, and blood and bone marrow were collected for flow cytometry analysis.

**Organ and tissue processing.** Venous blood was collected via right ventricle puncture into EDTA-coated tubes to prevent coagulation. Red blood cells were lysed with Lysing Buffer (BD Biosciences) for 5 minutes at room temperature. The cell suspension was washed twice with FACS buffer (PBS, 2% FBS, 0.1% EDTA) by centrifugation at 400 × g for 5 minutes at 4°C, discarding the supernatant each time. Cells were resuspended in 200 μl FACS buffer and stored on ice for further experiment; Bone marrow was isolated from mouse femurs using short time high-speed centrifugation. Red blood cells were lysed with lysis buffer for 30 seconds at room temperature. Cells were washed once with FACS buffer by centrifugation at 400 × g for 5 minutes at 4°C, supernatant discarded, and resuspended in 400 μl FACS buffer, kept on ice for analysis. Spleens were mechanically homogenized through a 70 μm cell strainer to yield a single-cell suspension. Cells were pelleted at 400 × g for 5 minutes at 4°C, followed by red blood cell lysis with lysis buffer for 2 minutes at room temperature. The suspension was washed once with FACS buffer at 400 × g for 5 minutes at 4°C, supernatant discarded, and cells resuspended in 800 μl FACS buffer, stored on ice for subsequent experiments.

**Flow cytometry analysis.** Cell suspensions from Bone marrow, Blood and Spleen were incubated with the flow cytometry antibodies and analysed using 5-lasers BD Fortessa X-20 (BD) and data was subsequently analyzed with FlowJo software (Tree Star). Quantification of cell numbers was done using count beads (CountBright; Life Technologies) following manufacturer’s instructions. After exclusion of cell doublets and dead cells with BD Fixable Viability Stain 700 (BD Biosciences), immature neutrophils were identified as (Lin/CD115/Siglec-F)^-^Gr1^+^CD11b^+^CXCR4^lo^cKit^lo^Ly6G^+^CD101^-^) and mature neutrophils were identified as (Lin/CD115/Siglec-F)^-^Gr1^+^CD11b^+^CXCR4^lo^cKit^lo^Ly6G^+^CD101^+^). Representative gating strategies for individual flow cytometry experiments are provided in Supplementary materials and used antibodies are shown as following:

| Company | Antibody | Clone | Reference | Dilution |
| --- | --- | --- | --- | --- |
| BioLegend | BV 510 anti-mouse CD117 | 2B8 | 105839 | 1:200 |
| BioLegend | BV 605 anti-mouse CD115 | AFS98 | 135517 | 1:200 |
| BioLegend | BV 711 anti-mouse Ly6G | 1A8 | 127643 | 1:200 |
| BioLegend | BV 785 anti-mouse/human CD11b | M1/70 | 101243 | 1:200 |
| BD | APC-Cy7 anti-mouse CD45 | 30-F11 | 557659 | 1:200 |
| Invitrogen | PE-Cy7 anti-mouse CD101 | Moushi101 | 25-1011-82 | 1:200 |
| BioLegend | AF488 anti-mouse Ly6C | HK1.4 | 128022 | 1:200 |
| BD | PerCP/Cy5.5 rat anti-mouse Gr-1 | RB6-8C5 | 552093 | 1:400 |
| BD | APC anti-mouse CD19 | 1D3 | 550992 | 1:200 |
| Invitrogen | APC anti-mouse NK1.1 | PK136 | 17-5941-82 | 1:200 |
| BioLegend | APC anti-mouse CD90.2 | 53-2.1 | 140311 | 1:200 |
| BD | Fixable viability stain 700 | / | 564997 | 1:1000 |
| BioLegend | BV 421 anti-mouse F4/80 | BM8 | 123137 | 1:200 |
| BD | BV 421 anti-mouse Siglec-F | E50-2440 | 562681 | 1:200 |
| BD | BUV 737 anti-mouse CD11b | M1/70 | 612800 | 1:200 |
| BD | BUV 395 anti-mouse CD4 | GK1.5 | 565974 | 1:200 |
| BD | BUV 496 anti-mouse CD45 | 30-F11 | 749889 | 1:200 |
| BioLegend | BV 510 anti-mouse TCR $\beta$ chain | H57-597 | 109233 | 1:200 |
| BioLegend | BV 605 anti-mouse CD11c | N418 | 117334 | 1:200 |
| BioLegend | BV 650 anti-mouse CD8a | 53-6.7 | 100742 | 1:200 |
| BD | BV 711 anti-mouse I-A/I-E | M5/114.15.2 | 563414 | 1:200 |
| BioLegend | BV 785 anti-mouse Ly6C | HK1.4 | 128041 | 1:200 |
| BioLegend | AF488 anti-mouse Ly6G | 1A8 | 127625 | 1:200 |
| BioLegend | PerCP/Cy5.5 rat anti-mouse B220 | RA3-6B2 | 103236 | 1:200 |
| Invitrogen | APC anti-mouse NK1.1 | PK136 | 17-5941-82 | 1:200 |

### Cytospin and Wright-Giemsa staining

To examine their distinct morphological features, Wright-Giemsa staining was performed as previously described^7^. Briefly, sorted CD101^-^ / CD101^+^ / tdTomato^-^ / tdTomato^+^ subsets (5×10^4^ cells each) were spun onto glass slides using Epredia Cytospin 4 Cytocentrifuge (Thermofisher Scientific) and air-dried. The slide was stained using Wright-Giemsa Quick Stain Kit (abs9293, Absin) following manufacturer’s instructions. 20 μl Giemsa stain Reagent A was added directly onto the slide and incubated at room temperature for 2 min. Next, 40 μl Giemsa stain reagent B was added directly onto the slide with Reagent A and incubated at room temperature for 10 min. Then, the staining reagents were removed under running water and air-dried. Images were acquired by Inverted Fluorescence Microscope (Leica, DM2500) equipped with a 100x oil immersion objective, and image brightness and cell diameter measurement were processed with ImageJ.

### Oxidative Burst Assay

Oxidative burst activity was measured as a function of their ability to generate superoxide anion and hydrogen peroxide, as detected by oxidation of dihydrorhodamine (DHR) (D23806, Invitrogen)^7^. In brief, sorted Ly6G^+^ cells (5×10^4^ cells each) were incubated with DHR for 30 min at 37°C in RPMI (supplemented with 10% FBS, 1% penicillin/streptomycin and 1% L-glutamine) and subjected to 100 nM 12-Myristate 13-Acetate (PMA) (Sigma-Aldrich) for 30 min at 37°C. Cells were then washed with 1xPBS followed by immediate data acquisition using flow cytometry.

***In vitro* cell culture.** Sorted cells (5 x 10^4^ for each subset) were plated onto 24-well plate and cultured in Iscove’s modified Dulbecco’s medium (Sigma) with 25mM HEPES and L-Glutamine (Chemtron) containing 10% (vol/vol) FBS, 1 mM sodium pyruvate, penicillin (100 U/mL) and streptomycin (100 ug/mL) at 37 ℃, 5% CO_2_. Cells were then harvested at 0, 1, 4, 24, 48, and 72 hours post-culture and analyzed for cell death and apoptosis using the Dead Cell Apoptosis Kit (Invitrogen) according to the manufacturer’s instructions.

**Neutrophil migration assay.** The neutrophil migration assay was performed using Corning® 6.5 mm Transwell® with 5.0 µm Pore Polycarbonate Membrane Insert as described earlier^2^. In brief, sorted cells (1 x 10^5^ for each subset) were applied in triplicates in 200 μl to the surface of membrane insert. Correspondingly, a 500 μl solution of 10^−6^ M fMLP (Sigma Aldricht) was added to the lower chamber. As a 100% migration control, 1 x 10^5^ cells for each subset were plated in duplicates in wells without inserts. The migration system was incubated at 37 ℃ with 5% CO_2_ for 4 hours. Cells that migrated to the lower chamber were harvested. Neutrophil migration rate was calculated as: (cell numbers below the insert / 100% migration control) x 100%.

**Quantification and statistical analysis.** Statistical analyses were performed using GraphPad Prism software. The Mann-Whitney U test was used for comparisons between two groups. For comparisons involving three or more groups, the Kruskal-Wallis test with Dunn’s multiple comparison correction was applied. All data are presented as means ± standard deviations, with replicate numbers specified in the figure legends.

## ACKNOWLEDGEMENTS

We would like to thank Rui He for the outstanding technical support and the animal caretakers from Shanghai Jiao Tong University School of Medicine for the excellent care provided to the animals. We are also grateful to Dongcheng Gong and Dongliang Xu from the technical platform of Shanghai Immune Therapy Institute, Shanghai Jiao Tong University School of Medicine Affiliated Renji Hospital, for their valuable technical assistance. This work was supported by funding to Lai Guan Ng from the Chinese Ministry of Science and Technology (grant number 2023YFC2306300) and the Research Fund for International Senior Scientist (grant number W2431020), and the Shanghai Science and Technology Development Funds (grant number 24PJA072), China Postdoctoral Science Foundation (2024M762041), Basic-Clinical Collaborative Innovation Project from Shanghai Immune Therapy Institute.

---

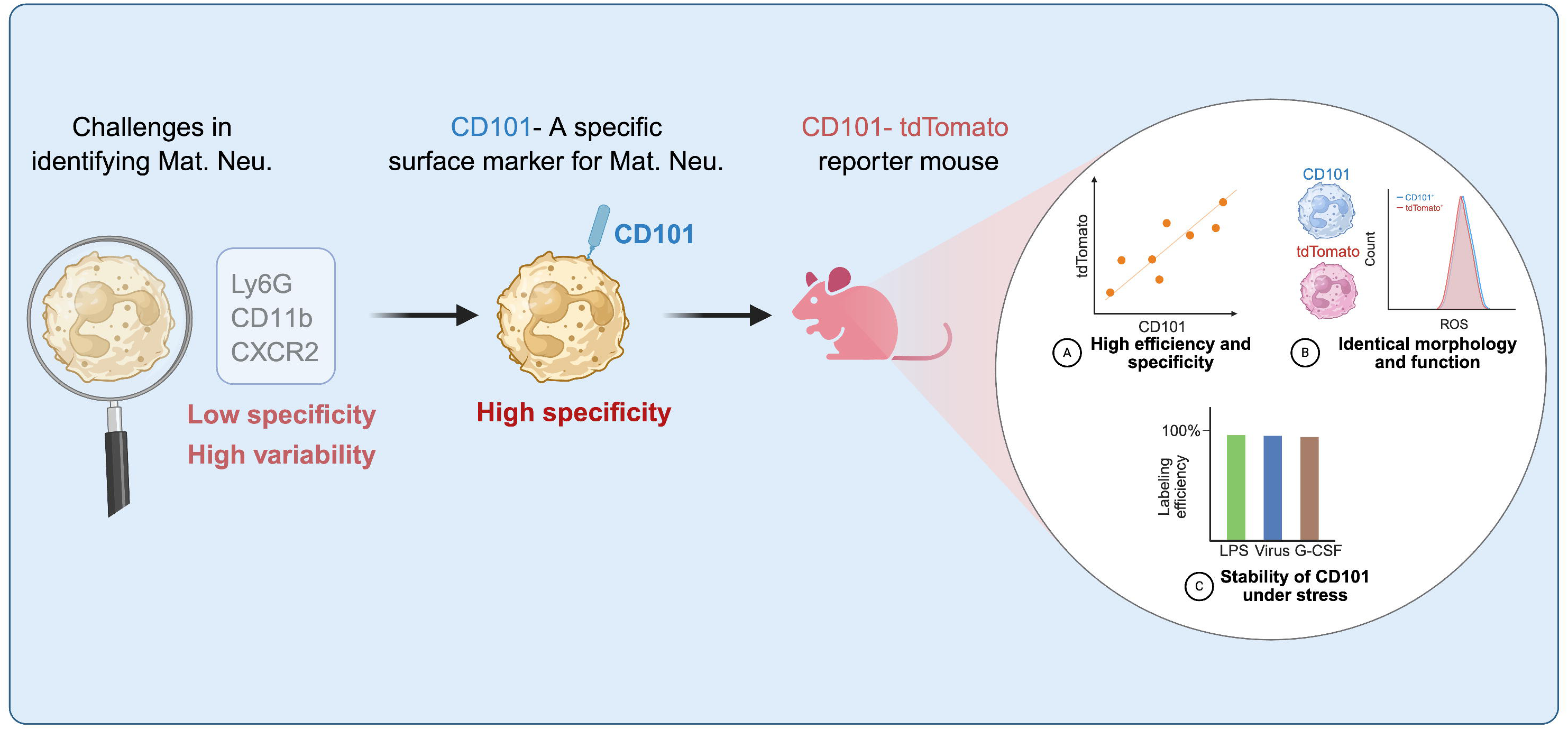

