## Supplementary Figures for "A Genetic Tool for Specific Tracking of Mature Neutrophils"

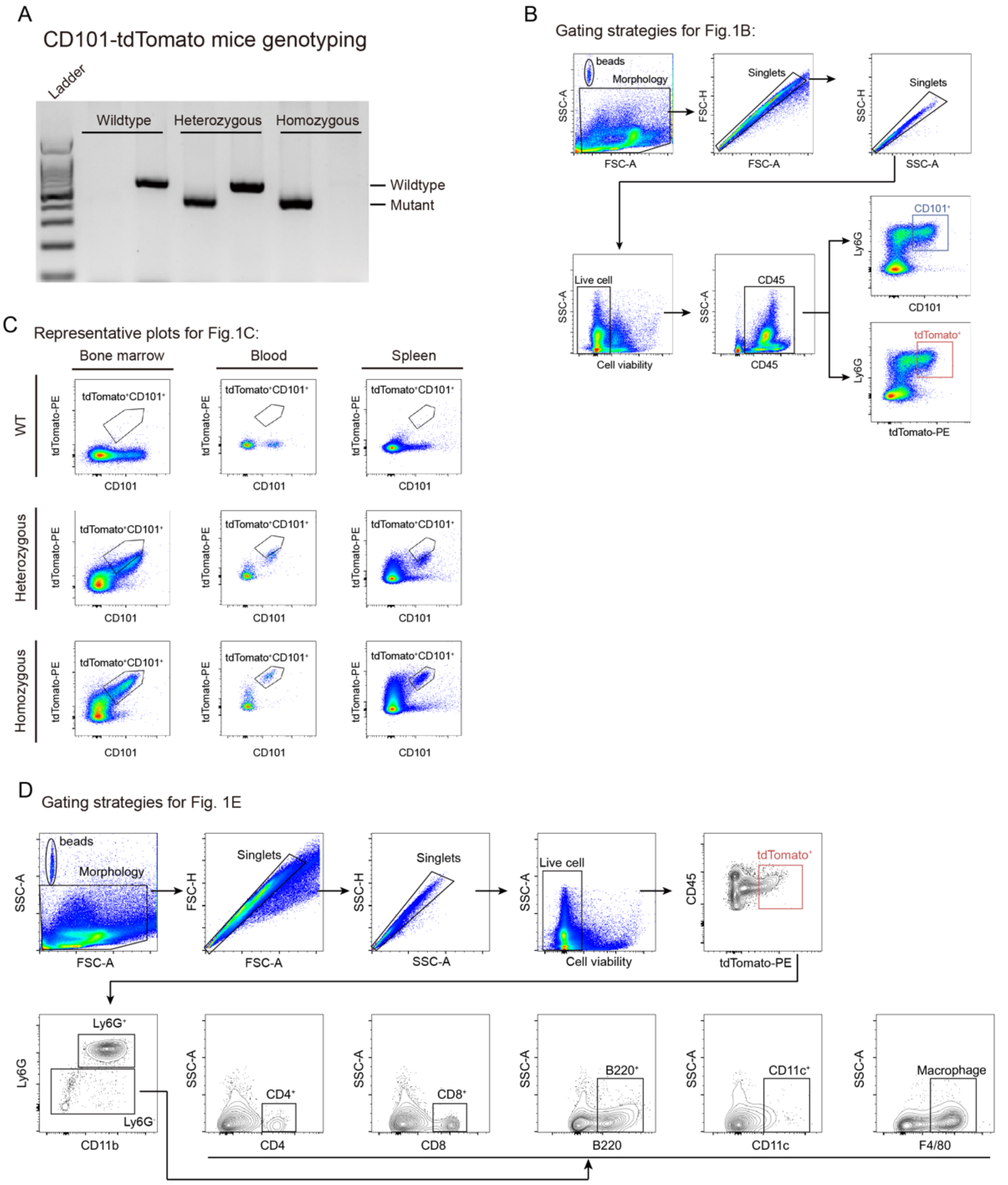


**Supplementary Figure 1. A.** Representative images of genotyping for wild-type (WT), heterozygous, and homozygous CD101-tdTomato mice, depicting the PCR-based detection of the DNA extracted from toe biopsies; **B.** Representative gating strategies of CD101$^{+}$ and tdTomato$^{+}$ mature neutrophils in BM for data showing in Fig. 1 B; **C.** Representative flow cytometry plots of tdTomato$^{+}$ against CD101$^{+}$ in WT, heterozygous, and homozygous CD101-tdTomato mice in peripheral organs for data showing in Fig. 1 C; **D.** Representative gating strategies to define tdTomato signal in general immune cells for data showing in Fig. 1 E.

Gating strategies for Fig.2A

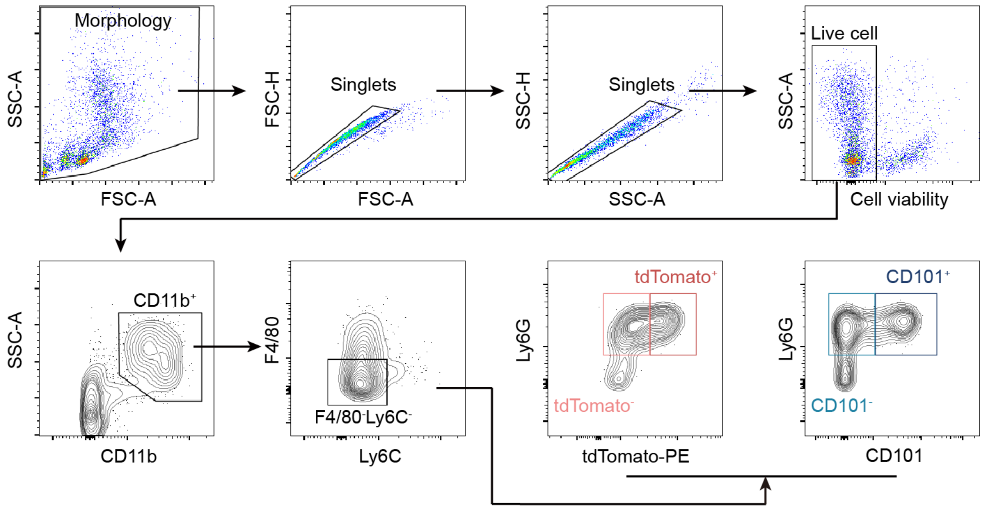


**Supplementary Figure 2. A.** Representative sorting strategies of mature / immature neutrophils sorted for nucleus morphology comparison for data showing in Fig. 2A;.

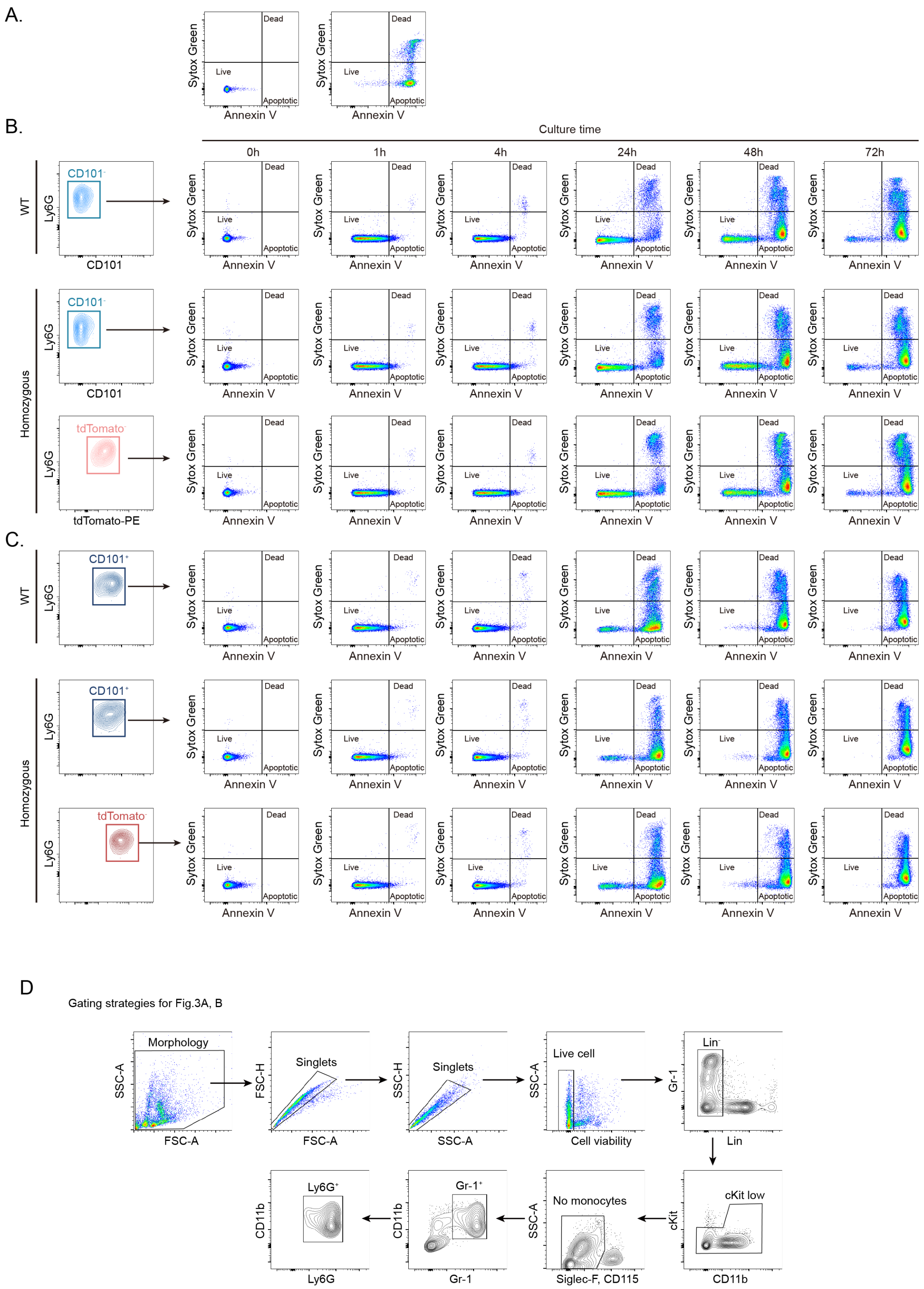


**Supplementary Figure 3. A.** Representative images of live cells, apoptotic cells and dead cells in negative control (left) and $H_2O_2$ treated cells; **B.** Representative flow cytometry plots showing the proportions of live, apoptotic, and dead cells among sorted CD101$^-$ and tdTomato$^-$ immature neutrophils from BM of WT and homozygous CD101-tdTomato mice, assessed at 0, 1, 4, 24, 48, and 72 hours post-culture for data showing in Fig. 3A; **C.** Representative flow cytometry plots depicting the proportions of live, apoptotic, and dead cells among sorted

CD101$^+$ and tdTomato$^+$ mature neutrophils from BM of WT and homozygous CD101-tdTomato mice, evaluated at 0, 1, 4, 24, 48, and 72 hours post-culture for data showing in Fig.3B. **D.** Representative sorting strategies for Ly6G$^+$ neutrophils sorted for ROS production for data showing in Fig. 3 C, D

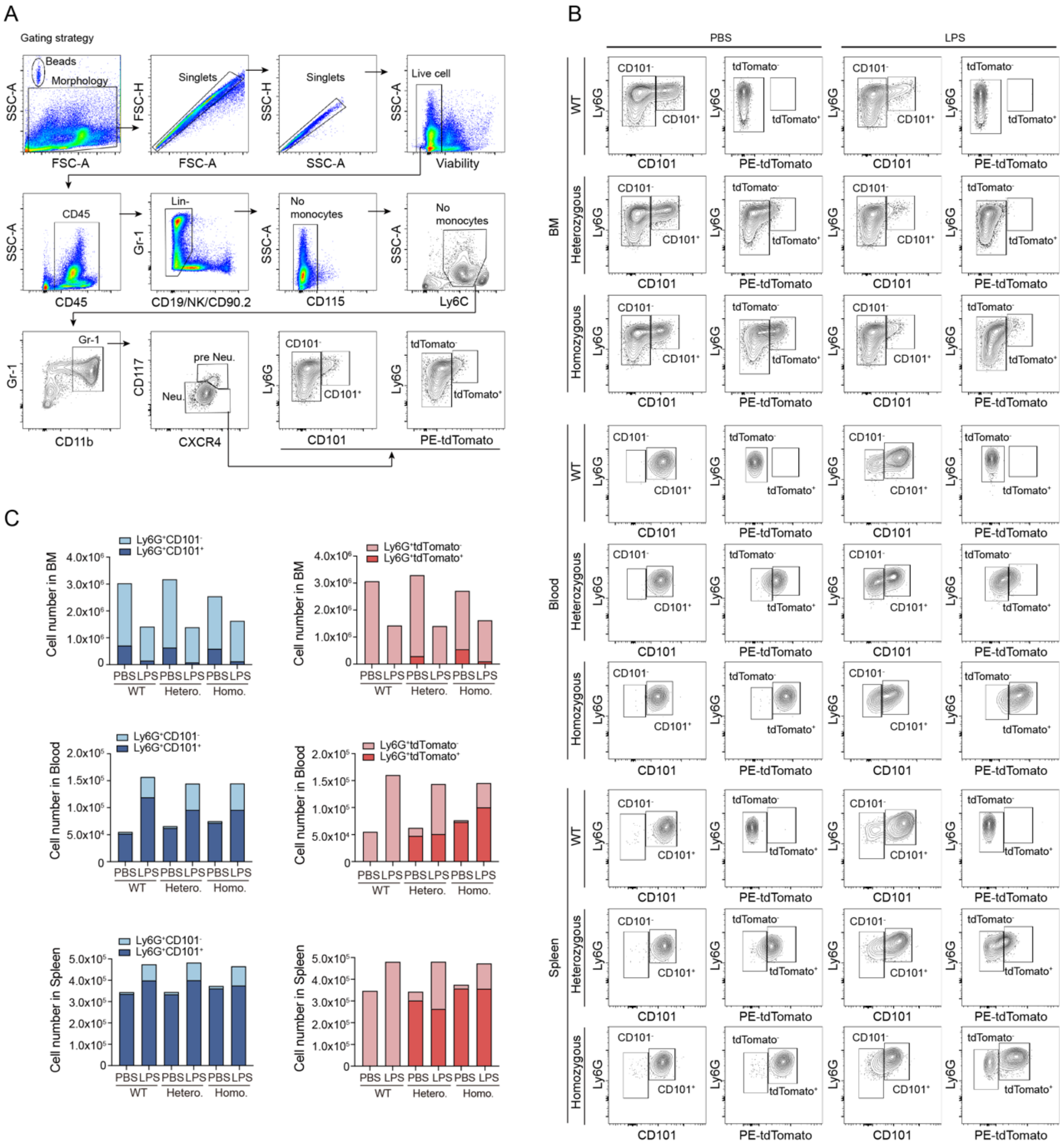


**Supplementary Figure 4. A.** Representative gating strategies of CD101$^{+}$ and tdTomato$^{+}$ mature neutrophils changes in BM after LPS challenge for data showing in Fig. 4 A; **B.** Representative flow cytometry plots illustrating the changes in CD101$^{+}$ and tdTomato$^{+}$ neutrophil populations in BM peripheral blood, and spleen from WT, heterozygous, and homozygous CD101-tdTomato mice under control conditions vs LPS challenge; **C.** Statistical analysis quantifying the proportion of mature and immature neutrophils in BM, peripheral blood, and spleen of WT, heterozygous, and homozygous mice under control and LPS-treated conditions (*n* = 8 per group).

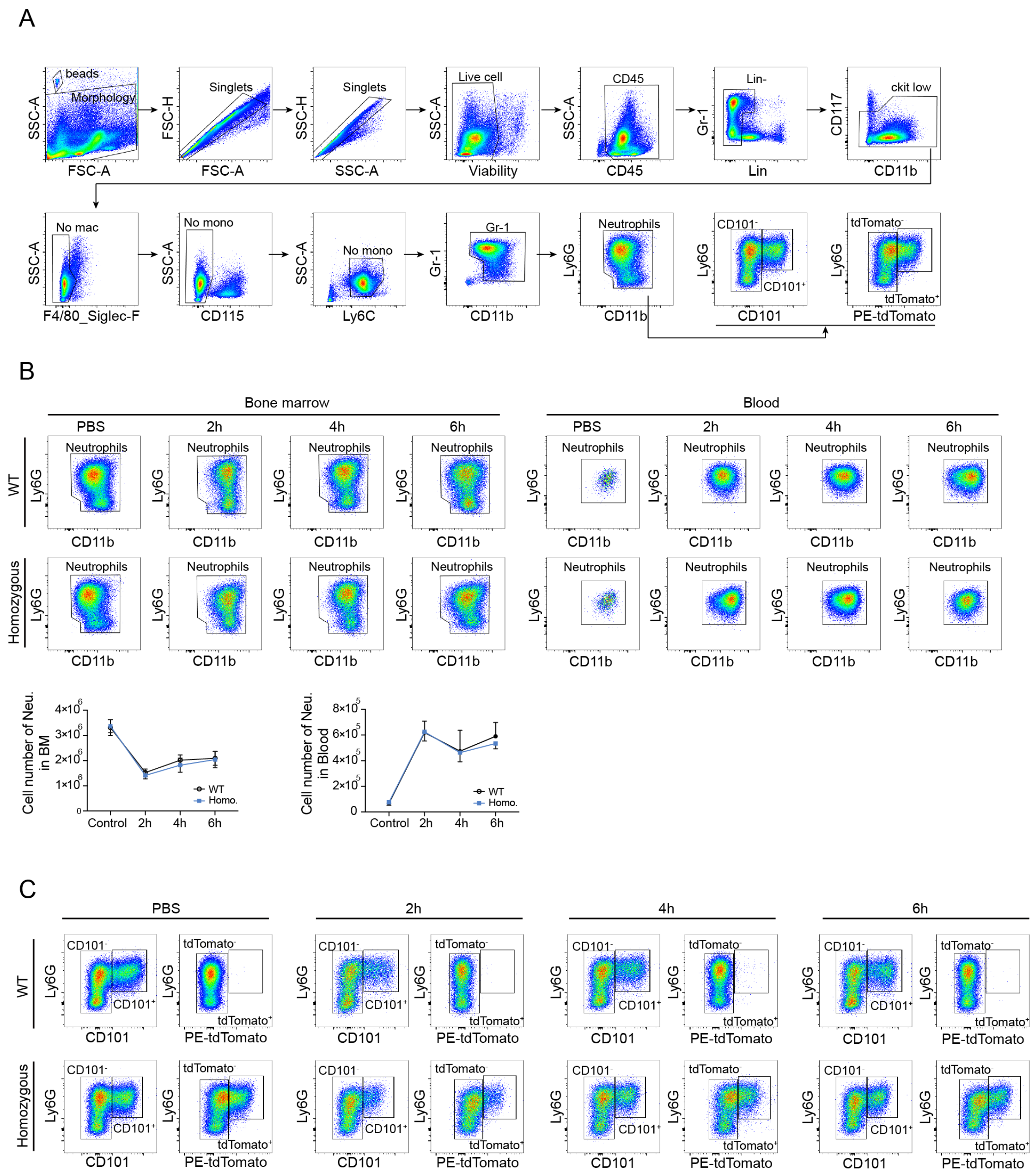


**Supplementary Figure 5. A.** Representative gating strategies of neutrophil dynamics in BM after G-CSF injection; **B.** Representative flow cytometry plots (upper) and corresponding quantification (lower) showing changes in total neutrophil populations in BM and peripheral blood of WT and homozygous CD101-tdTomato mice at 2, 4, and 6 hours following G-CSF injection (*n* = 5 per group); **C.** Representative flow cytometry plots illustrating dynamics in mature (CD101$^+$ and tdTomato$^+$) and immature (CD101$^-$ and tdTomato$^-$) neutrophil populations in BM and peripheral blood of WT and homozygous mice following G-CSF injection for data showing in Fig. 4 D

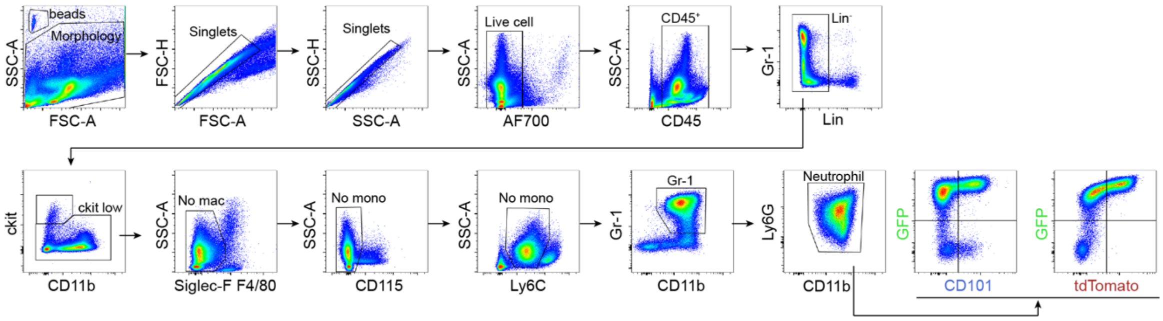


**Supplementary Figure 6. A.** Representative gating strategies of distinguishing mature neutrophils by using CD101-tdTomato Lysozyme GFP mice;
